# Phylogenetic analysis of human parainfluenza type 3 virus strains responsible for the outbreak during the COVID-19 pandemic in Seoul, South Korea

**DOI:** 10.1101/2022.03.15.484550

**Authors:** Ha Nui Kim, Soo-Young Yoon, Chae Seung Lim, Chang Kyu Lee, Jung Yoon

## Abstract

**Background:** Human parainfluenza virus 3 (HPIV3) is a major respiratory pathogen that causes acute respiratory infections in infants and children. Since September 2021, an out-of-season HPIV3 rebound has been noted in Korea. The objective of this study was to analyze the molecular characteristics of the HPIV3 strains responsible for the outbreak in Seoul, South Korea.

**Methods:** A total of 61 HPIV3-positive nasopharyngeal swab specimens were collected between October and November 2021. Using 33 HPIV3-positive specimens, partial nucleotide sequences of the HPIV3 hemagglutinin-neuraminidase (*HN*) gene were aligned with previously published *HN* gene sequences for phylogenetic and genetic distance (p-distance) analyses.

**Results:** **P**hylogenetic tree revealed that all Seoul HPIV3 strains grouped within the phylogenetic subcluster C3. However, these strains formed a unique cluster that branched separately from the C3a lineage. This cluster showed 99% bootstrap support with a p-distance < 0.001. Genetic distances within the other C3 lineages ranged from 0.013 (C3a) to 0.023 (C3c). Deduced amino acid sequences of the *HN* gene revealed four protein substitutions in Seoul HPIV3 strains that have rarely been observed in other reference strains: A22T, K31N, G387S, and E514K.

**Conclusions:** **P**hylogenetic analysis of Seoul HPIV3 strains revealed that the strain belonged to a separate cluster within subcluster C3. Genetic distances among strains within subcluster C3 suggest the emergence of a new genetic lineage. The emergence of a new genetic lineage could pose a potential risk of a new epidemic. Further monitoring of the circulating HPIV3 strains is needed to understand the importance of newly discovered mutations.

## 1. Introduction

Human parainfluenza virus (HPIV) is an important causative pathogen of acute lower respiratory tract infections (LRTI). HPIV usually causes mild respiratory infections in adults, which are indistinguishable from the common cold; however, it is often responsible for hospitalization in children under 5 years of age or children with underlying conditions. The incidence of LRTI caused by HPIV in young children is known to be surpassed only by human respiratory syncytial virus (RSV) (1). HPIVs have been classified into four types: 1, 2, 3, and 4. HPIV3, which belongs to the genus *Respirovirus* of the family *Paramyxoviridae*, is the most prevalent type among HPIVs and is commonly associated with LRTIs (2, 3). HPIV3 infections show clear seasonality and are prevalent during late spring and summer (4).

Since the coronavirus disease 2019 (COVID-19) pandemic, non-pharmaceutical interventions (NPIs) have been widely implemented to reduce transmission (5). These interventions against COVID-19 likely reduced the activity of other respiratory viral infections, and a substantial decrease in the incidence of respiratory viral infections other than COVID-19 has been reported (6, 7). Interestingly, in late 2020, a delayed outbreak of RSV was reported in some countries (8). In Korea, an out-of-season HPIV3 outbreak has been reported during the COVID-19 pandemic in September 2021. From the 39^th^ to 42^nd^ week of 2021, an unprecedented high positivity rate of 41.6% was noted (9). Moreover, the HPIV3 epidemic differed in that the HPIV3 detection was the most prevalent among infants and children, and other types of HPIV were not detected (9, 10).

HPIV3 is an enveloped, negative-sense, single-stranded RNA virus, with 15,462 nucleotides that encode six genes and eight proteins (3). The major antigens responsible for infecting the host cells are its two surface glycoproteins; the hemagglutinin-neuraminidase (HN) and fusion (F) proteins. The HN protein regulates virus-host interactions and has both hemagglutinin and neuraminidase activity (11). In addition, the HN protein affects the F protein’s function in mediating membrane fusion with host cells (12). Because of its antigenicity and variability, the *HN* gene has been the preferred target gene region for the phylogenetic analysis of HPIV3 (13). HPIV3s are classified into three clusters (clusters A–C) according to the characteristics of the *HN* gene (11), and previous studies have revealed that cluster C can be subdivided into five subclusters (C1–C5) based on the nucleotide sequence of the *HN* gene (14). Monitoring the amino acid variations occurring in the *HN* gene may help identify the effects of these substitutions on HN protein function and viral evolution (15). In this study, we aimed to reveal the molecular characteristics of the HPIV3 strains responsible for an out-of-season outbreak in Seoul, South Korea and understand the molecular evolution of HPIV3 detected during the COVID-19 pandemic.

## 2. Materials and methods

### 2.1. Sample and clinical data collection

A total of 307 nasopharyngeal swab (NPS) specimens were collected from patients presenting with respiratory symptoms at the Korea University, Guro Hospital between October and November 2021. Among 307 NPS specimens, 61 specimens were HPIV3-positive in a multiplex real-time (RT) polymerase chain reaction (PCR) assay (Anyplex II RV16 Detection Kit, Seegene, Seoul, Korea), and were included in our study. Clinical data, including patient demographic characteristics, were collected from the clinical records of the hospital. This study was approved by the Institutional Review Board (IRB) of the Korea University Guro Hospital (IRB number: 2022GR0003). Partial sequencing of the *HN* gene was performed using 37 HPIV3-positive samples.

### 2.2 RNA extraction and RT-PCR for the analysis of partial sequence of the *HN* gene

Viral RNA was extracted from the NPS samples using the nucleic acid extraction platform Microlab STARlet (Hamilton, Reno, Nevada, USA), and cDNA was synthesized using the RevertAid First Strand cDNA Synthesis kit (Thermo Fisher Scientific, Waltham, MA, USA) according to the manufacturer’s instructions. For amplification of the partial nucleotide sequence of *HN* gene, previously described target gene-specific primer pairs (13) and Dr. MAX DNA polymerase (Doctor Protein Inc., Seoul, Korea) were used. PCR conditions were as follows: initial denaturation at 95°C for 5 minutes; followed by 35 cycles of 95°C for 30 s, primer annealing at 60°C for 30 s, and extension at 72°C for 1 minute, and a final extension step at 72°C for 7 minutes. The amplified PCR products were visualized by 1.5% agarose gel electrophoresis and purified using a Millipore filter plate MSNU030 (Millipore SAS, Molsheim, France). The purified PCR products were then subjected to Sanger sequencing using the BigDye Terminator v3.1 sequencing kit and a 3730xl automated sequencer (Applied Biosystems, Foster City, CA). The obtained sequencing data were analyzed using the Variant reporter computer software version 1.1 (Applied Biosystems, Foster City, CA, USA).

### 2.3. Phylogenetic analysis

For phylogenetic and evolutionary analyses, data for 83 reference strains were retrieved from GenBank (https://www.ncbi.nlm.nih.gov/genbank/). The strains representing each HPIV3 cluster were collected from various countries and over time from 1957 to 2015. Multiple sequence alignment was performed using the nucleotide sequences of the *HN* gene obtained in this study with those of the reference strains, using the MUSCLE algorithm (16) implemented in MEGA X software (17). A maximum likelihood (ML) tree was constructed under the best-fit TN93 + G substitution model, as predicted by MEGA X. The reliability of the branching order was assessed using 1,000 bootstrap replicates, and clades with values > 70% were considered significant.

In addition, a time-scaled phylogenetic tree was reconstructed using the Bayesian Markov chain Monte Carlo (MCMC) method using BEAST v2.6.6 (18). The best-fit substitution model was determined using ModelFinder with TIM+F+I+G4 based on the minimum Bayesian information criterion value (19). The best model among the six clock-tree model combinations, the relaxed clock log-normal and constant coalescent population, was chosen by path sampling and stepping stone, using the model-selection package implemented in BEAST. The MCMC chain length was 300,000,000 steps, which were sampled every 5,000 steps. The Tracer program v1.7.1 (20) was used to ensure that the effective sample size was greater than 200 for all the parameters. The maximum clade credibility tree information was summarized using TreeAnnotator v2.6.6 (21) after burning in 10% of the trees. The Bayesian MCMC phylogenetic tree was visualized using FigTree v1.4.4 (22). To analyze the phylogenetic distances within and between HPIV3 clusters, the genetic distances (p-distances) of HPIV3 clusters, including standard error (SE), were calculated using MEGA X with 1,000 replicates by the bootstrap method.

### 2.4. Amino acid sequence analysis

The amino acid sequences were obtained by translating the nucleotide sequences of the partial nucleotide sequence of the *HN* gene using the standard genetic code in MEGA X. The deduced amino acid sequences were aligned with those of previously published strains of each cluster, including the prototype Washington/1957 strain (GenBank accession number: M17641), as a reference. Mutations at each codon position were visualized using Unipro UGENE (23). The crystal structure of HN protein was visualized using PyMOL 2.0 (24), based on the previously published crystal structure of HN (PDB ID: 1V3B). *In silico* mutagenesis was performed using PyMOL 2.0, to assess the impact of mutations on the protein structure. Entropy analysis by Shannon entropy plot was performed using BioEdit, with a threshold value of 0.2 (25).

### 2.5. Selection pressure analysis

The selection pressure on the HN gene of HPIV3 was estimated using the Datamonkey server (http://www.datamonky.org/). The non-synonymous (*d*N) and synonymous (*d*S) substitution rates at each amino acid position were calculated using the Mixed Effects Model of Evolution (MEME) and Fast, Unconstrained Bayesian AppRoximation (FUBAR) methods for the estimation of positive and negative selection sites, respectively (26, 27). The evidence for positive selection (dN/dS > 1) and negative selection (dN/dS <1) were supported by a p-value threshold of 0.05 and posterior probability of 0.9, respectively.

## 3. Results

### 3.1. Patient characteristics and clinical information

Among 307 NPS specimens, 19.87% (n = 61/307) were positive for HPIV3 in the multiplex RT-PCR assay. Of the 61 HPIV3-positive samples that were included in this study, 32 (52.5%) belonged to male, and 29 (47.5%) belonged to female patients. The median age of the patients was 24 months, and most (70.5%) were infants and children aged below 7 years (Table 1). A diagnosis of upper respiratory tract infection (URTI) was common (67.2%). The manifestation as LRTI including pneumonia and bronchiolitis was observed in 34.4% and 6.6% of patients, respectively. The most frequent diagnosis in infants and children was URTI (83.7%, n = 36/43), followed by croup (25.6%, n = 11/43), bronchiolitis (9.3%, n = 4/43), and pneumonia (9.3%, n = 4/43).

**Table 1.**
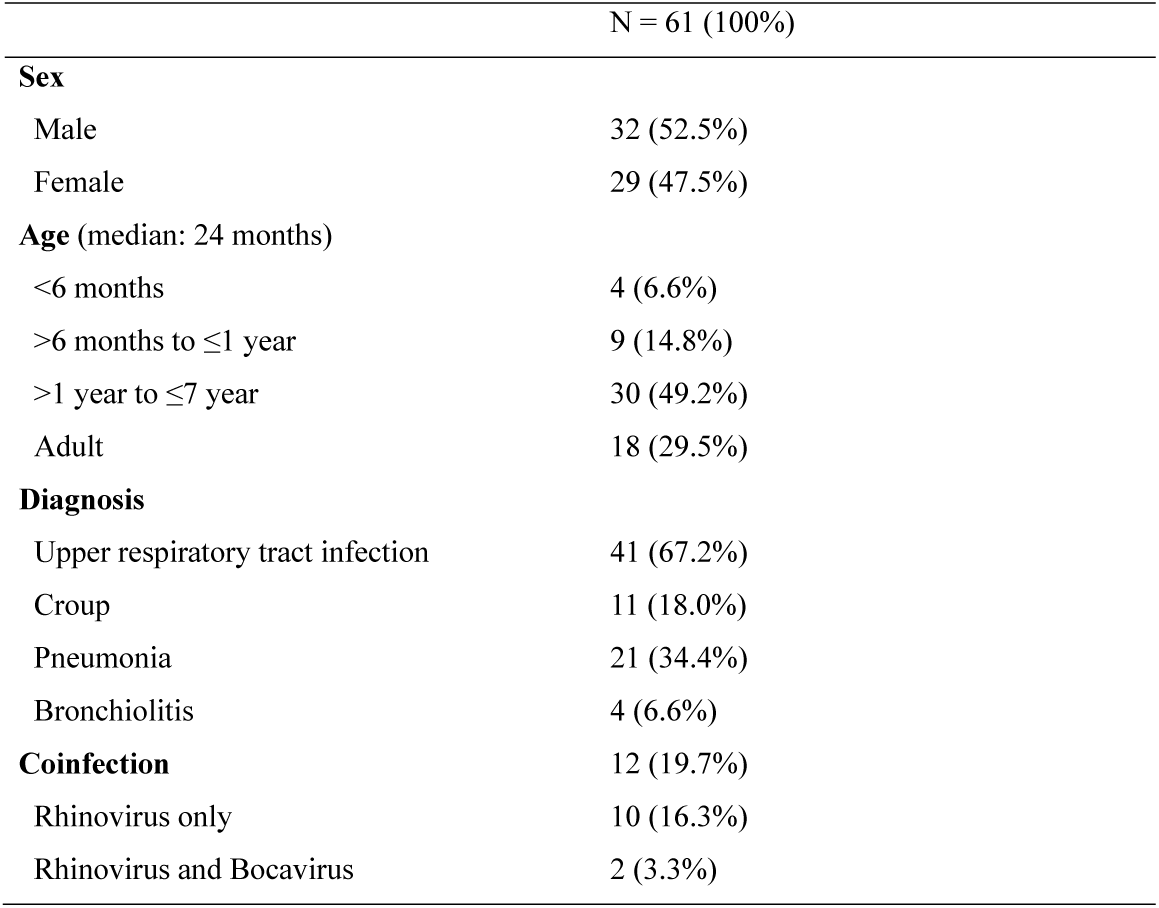
Characteristics and clinical information of patients whose samples were investigated in this study

### 3.2. Seasonality analysis

The HPIV3-positive cases reported at the Korea University Guro Hospital were investigated for seasonality analysis based on data in the last 5 years (2017–2021). A distinct seasonal prevalence of HPIV3 infection was observed during late spring and summer was observed until 2019, with the highest prevalence observed in May, ranging from 8.1% to 12.7%. However, the prevalence pattern in 2021 was different, with the emergence of an unexpected peak beginning in September, which extended into the winter, and a higher positivity rate was noted (23.9% in November) (Figure 1).

**Figure 1.**
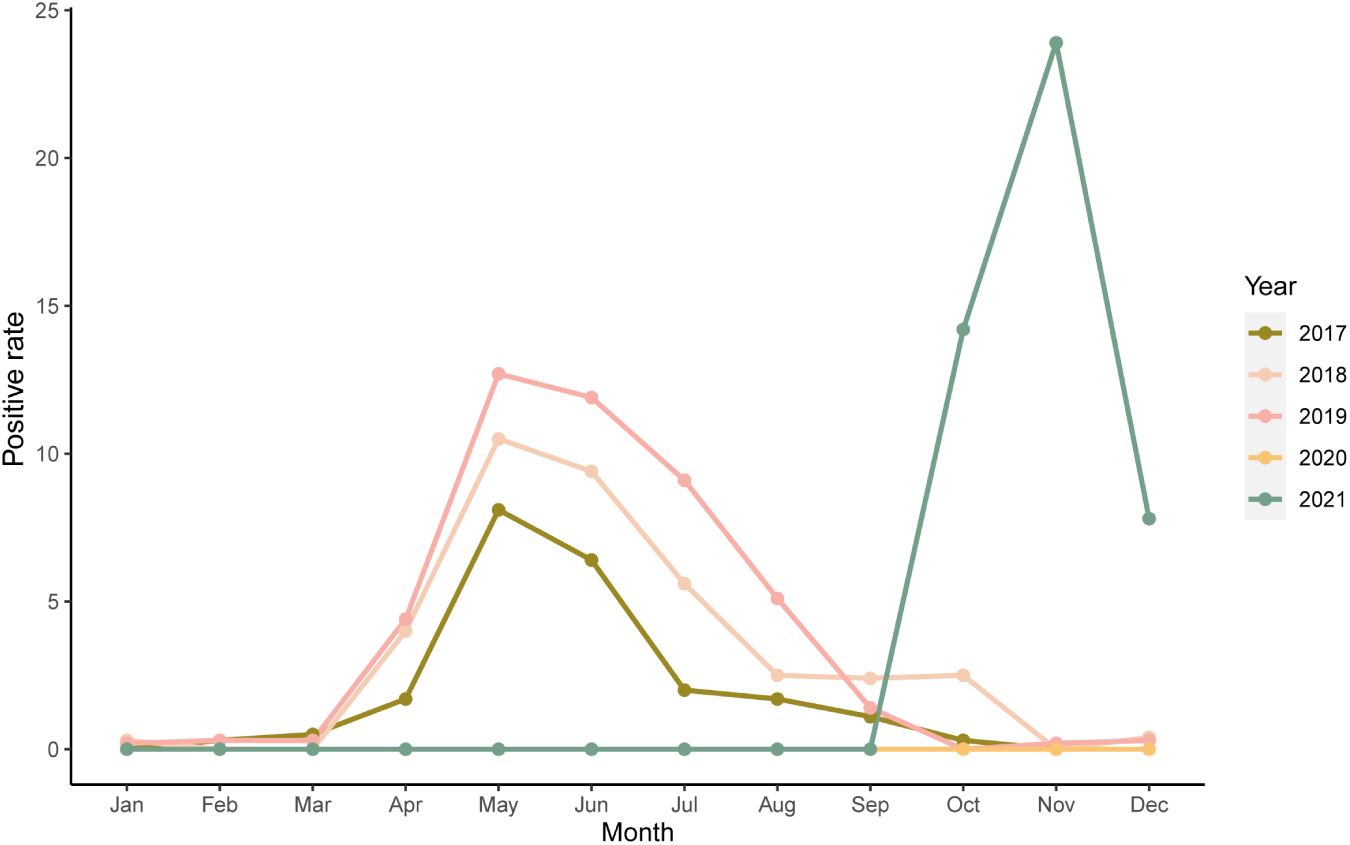
Seasonality pattern of human parainfluenza virus 3 (HPIV3)-positive cases per month between 2017 and 2021. The positivity rate was defined as the number of HPIV3 positive cases compared to the total number of multiplex real-time polymerase chain reaction tests.

### 3.3. Phylogenetic analysis

The 33 partial HN sequences obtained from the Seoul HPIV3 strains were aligned with the 88 reference sequences representing each cluster and genetic lineage. We could not obtain data for four samples; hence, those samples were excluded (Figure S1). This failure was likely due to the low viral load in each specimen. The ML tree showed that all Seoul HPIV3 strains formed a distinct cluster branching from C3a, with a high bootstrap value of 99% (Figure 2). The genetic distance within the Seoul HPIV3 group was 0.0004, whereas the genetic distance from the Seoul HPIV3 strains to other genetic lineages of subcluster C3 was 0.013 (C3a)–0.023 (C3c) (Table 2). The previously defined criteria for classification of clusters and subclusters were as follows: genetic distances ≥ 0.045, different clusters; distances in the range of 0.019–0.045, same clusters but different subclusters; 0.010–0.019, same subclusters but different genetic lineages (14, 28, 29). According to these criteria, the Seoul HPIV3 strains were considered a new genetic lineage within subcluster C3, and thus provisionally designated in this study as C3h.

**Table 2.**
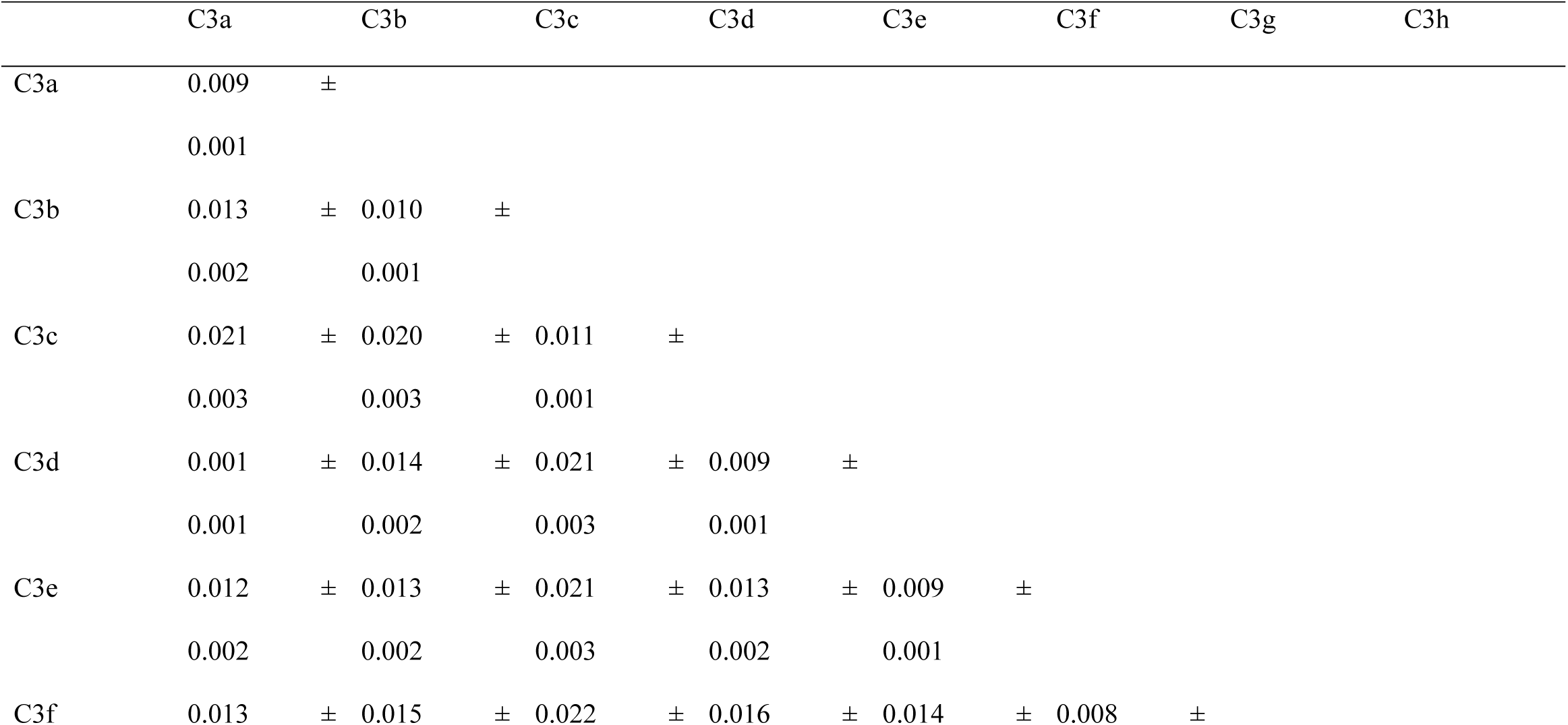
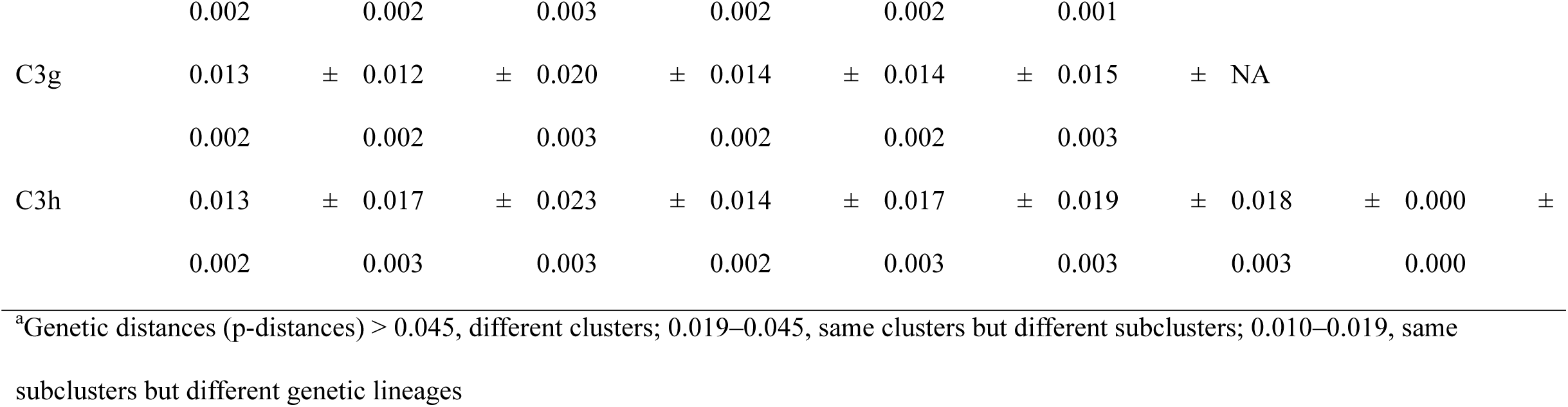
Genetic distances^a^ (mean ± SD) in human parainfluenza virus 3 (HPIV3) subcluster C3, including a new genetic lineage (Korean HPIV3, designated C3h)

**Figure 2.**
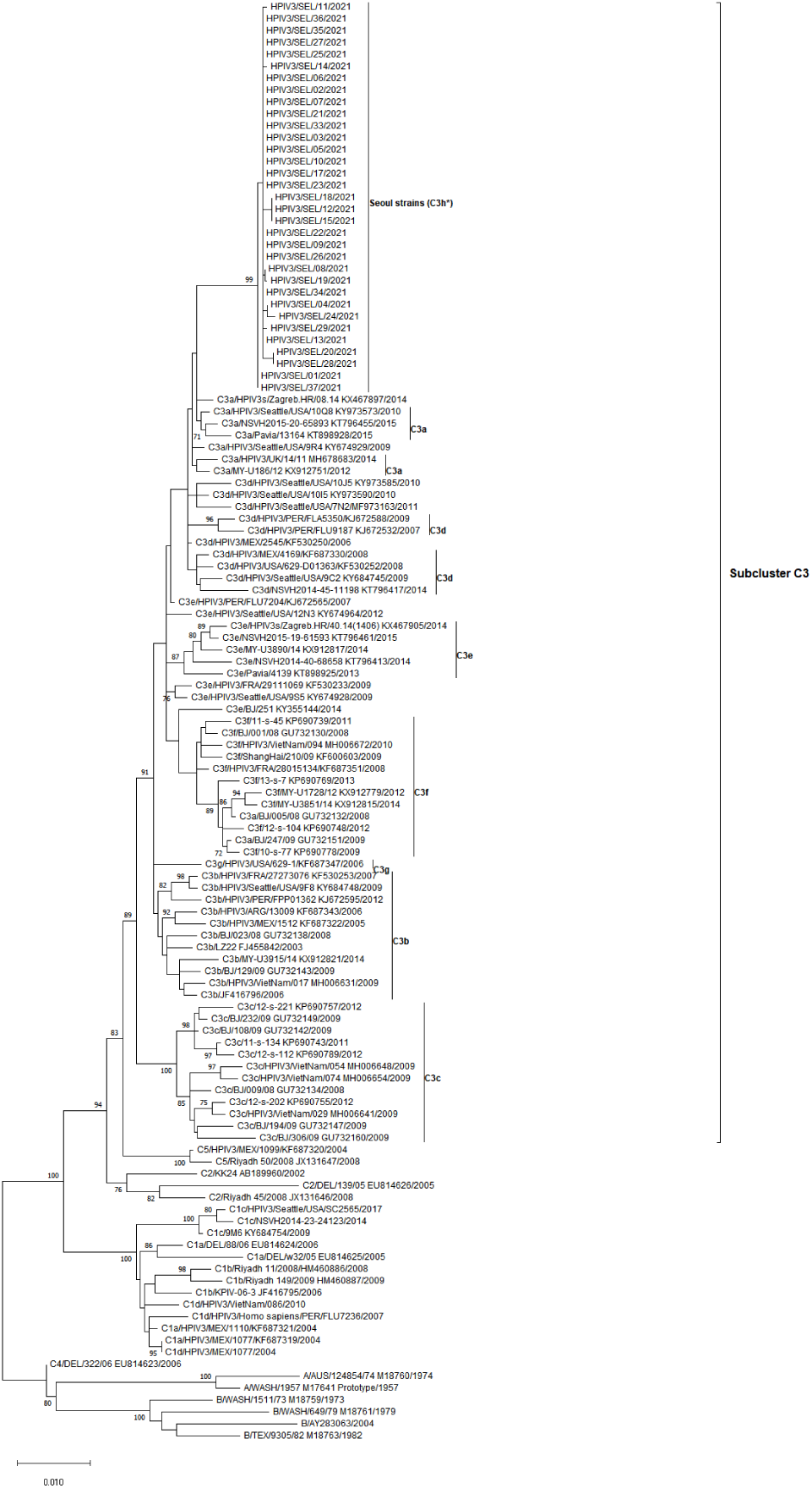
Maximum likelihood phylogenetic tree using partial nucleotide sequences of the *HN* gene of Seoul HPIV3 strains. The nucleotide sequences of the *HN* gene from strains representing their clusters and genetic lineages were aligned with those of the Seoul strains using MUSCLE implemented in MEGA X. The reliability of the tree was estimated using the bootstrap method with 1,000 replicates, and values above 70% are shown. Subcluster C3, genetic lineages of C3 (C3a – C3g), and Seoul strains (C3h) are indicated on the right-hand side of the phylogram.

Using the Bayesian MCMC method, the time-scaled phylogenetic tree of *HN* gene showed a similar result to that of the ML tree (Figure 3). The Seoul HPIV3 strains were grouped within subcluster C3 but formed a separate cluster distinct from C3a, with a posterior probability value of 0.95.

**Figure 3.**
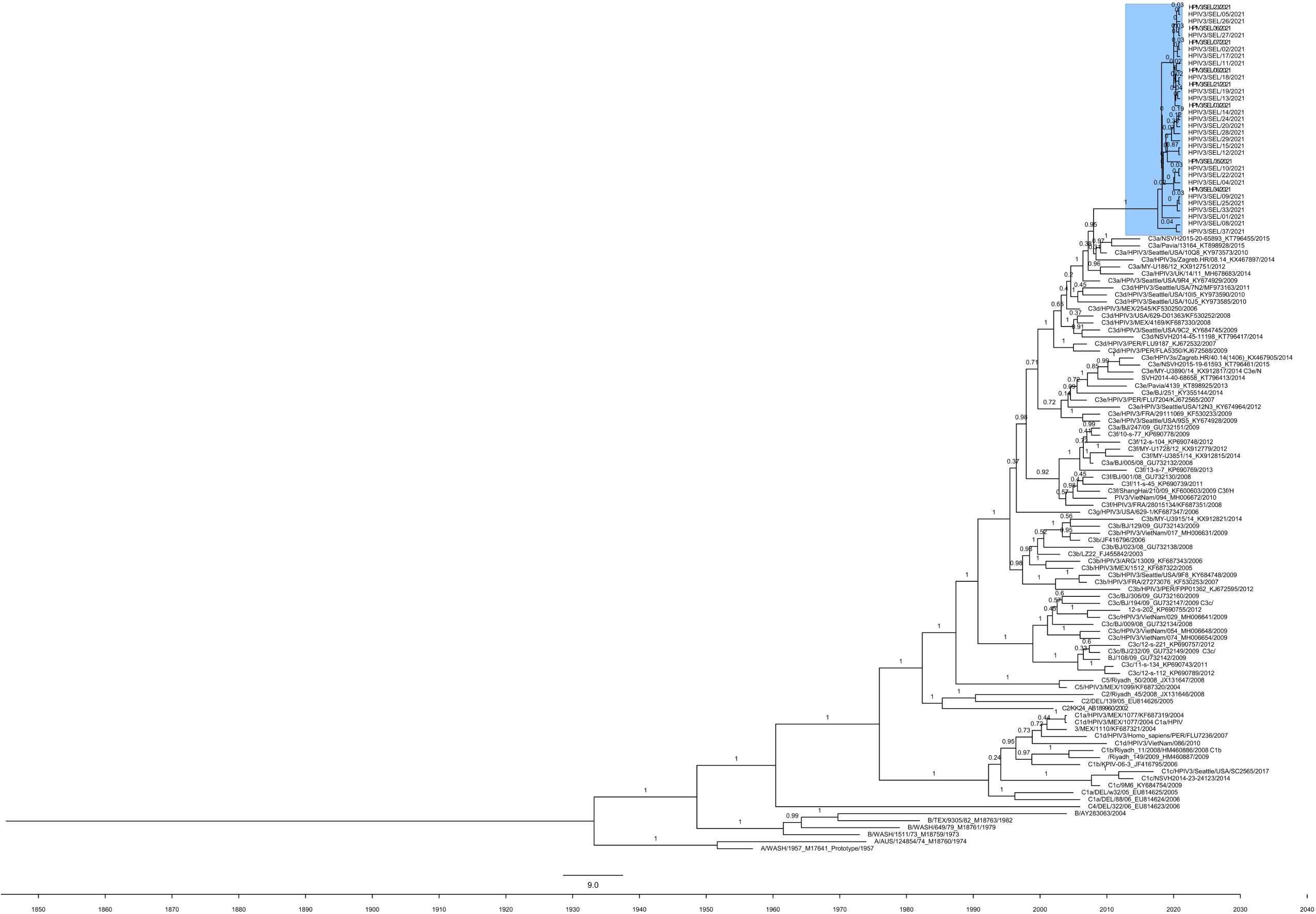
A time-scaled phylogenetic tree of the *HN* gene of Seoul HPIV3 strains was constructed using the Bayesian MCMC method. The Seoul HPIV3 strains are highlighted in a blue box.

### 3.4. Amino acid variations

All Seoul HPIV3 strains were compared with the prototype strain (GenBank accession number: M17641) in terms of their relative amino acid positions. Three common variations in cluster C (29) were observed in all the Seoul strains: H295Y, I391V, and D556N. Amino acid substitutions not frequently observed in reference strains were found in more than two strains in Seoul HPIV3 strains: A22T, K31N, G387S, and E514K. Four sequences from the Seoul HPIV3 strains harbored the E514K mutation, which was observed in only one of the reference strain (KJ672532, C3d). At the corresponding nucleotide position of 1540, a heterozygous mutation (G1540R) was observed in four Seoul HPIV3 strains (HPIV3/SEL21, 22, 26, and 29). This heterozygosity was supported by the forward and reverse amplicon reads. The locations of G387S and E514K in the HN crystal structure are shown in Figure 4.

**Figure 4.**
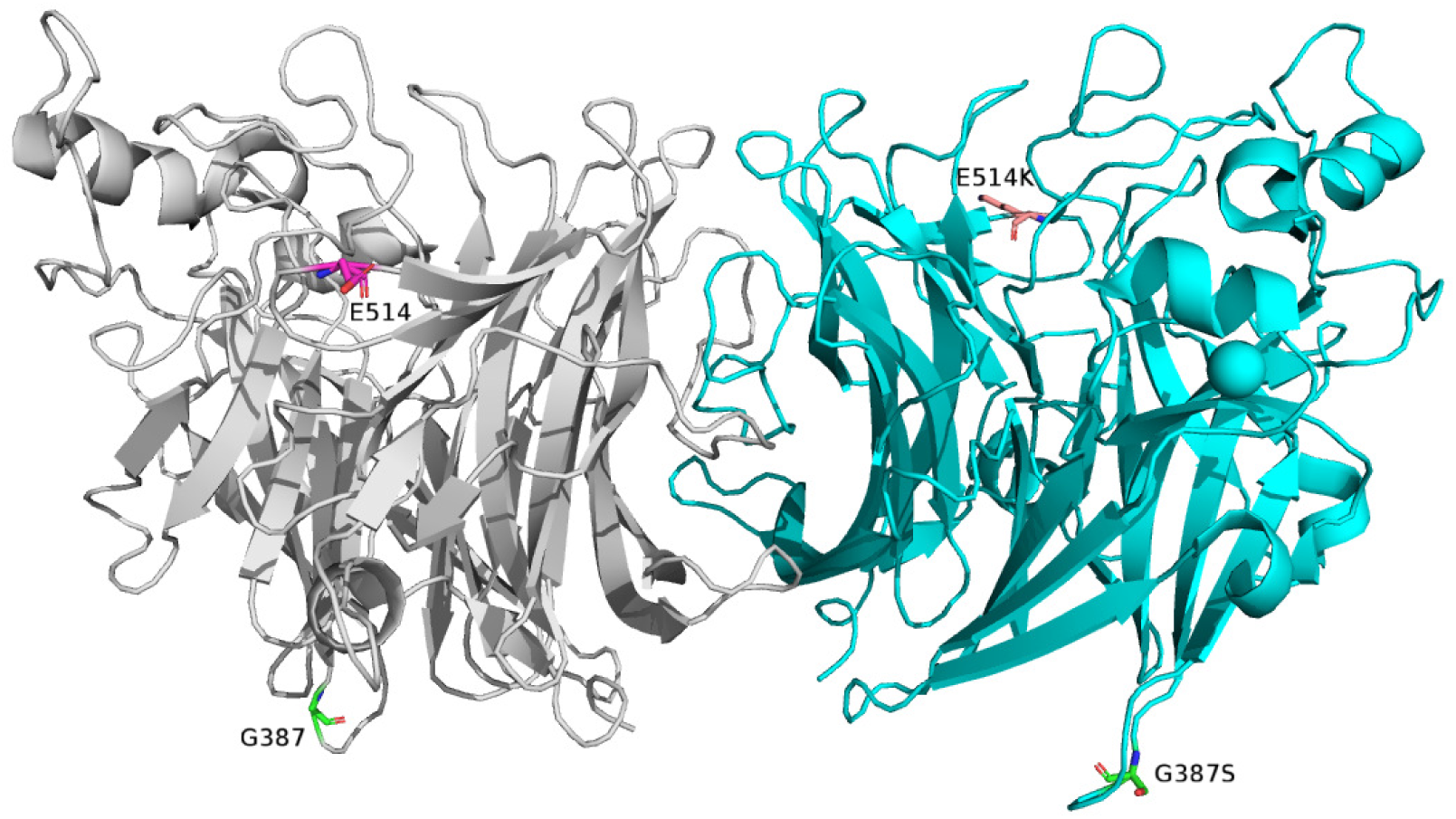
Molecular structure of the HPIV3 HN protein with monomers colored gray (left) and cyan (right). The structure was visualized using PyMol v2.0 (PDB no. 1V3B). The locations of G387 and E514 are highlighted as sticks in the left monomer, and G387S and E514K mutations after mutagenesis are indicated in the right monomer.

### 3.5. Positive and negative selection sites

We analyzed the positive and negative selection sites to identify selection pressure against the host using MEME and FUBAR. There were 15 positive selection sites. Many negative selection sites (total, 257 sites) were predicted, including A22T and G387S substitutions.

## 4. Discussion

As COVID-19 measures were relaxed, unexpected outbreaks due to infections by the suppressed respiratory viruses, so-called “out of season” epidemics, have been observed (30). The Korean government announced relaxation in COVID-19 restrictions on November 1, 2021, and discontinued “living with the COVID-19” scheme in December 2021. As the maximum number of infections due to HPIV3 strains were observed in November 2021, the anomalous occurrence of previously seasonal infections can be related to relaxed COVID-19 restrictions. The uncertainty with which respiratory virus infections can occur outside of their typical seasonality can pose a risk to the vulnerable populations. There are some concerns about the possibility of more severe epidemics when the respiratory viruses return because reduced natural exposure to these viruses can be lead to lowered population immunity (7).

In our study, compared to that before the COVID-19 pandemic, an almost 2-fold (ranging from 1.88-to 2.95-fold) increase in the peak positivity rate was detected. Unlike other HPIVs, HPIV3 mainly infects young infants and causes severe LRTI, such as bronchiolitis and pneumonia (1). Despite the high detection rate of HPIV3, especially in infants and children, compared to that reported in a previous study conducted in our institute (31), a lower frequency of pneumonia and bronchiolitis was observed. The most common manifestation of LRTI was croup.

According to recent studies from Argentina, Spain, and China, strains belonging to cluster C are predominantly prevalent (28, 32, 33). Phylogenetic analysis using the *HN* gene showed that all Seoul HPIV3 strains were classified as subcluster C3; however, they formed a distinct group. This finding was supported by both ML and time-scaled trees, with a bootstrap value of 99% and posterior probability value of 0.95. In addition, analysis of genetic distances within the subcluster C3 suggested the emergence of a newly subdivided genetic lineage C3h.

Similarly, the emergence of novel RSV lineages was reported after the COVID-19 pandemic in Australia (34). Previously prevalent RSV strains were mostly absent, and a novel lineage was dominant, with a significant reduction in G gene diversity. This major collapse in genetic diversity was also observed in influenza viruses, with dramatically decreased detection rates (35). The characteristics of HPIV3 identified in this study can be understood in this context.

Among the six proteins encoded by the HPIV3 genome, HN and F proteins are present in the viral lipid membrane (36). HN protein is a homotetrameric type II integral membrane protein consisting of a cytoplasmic domain, transmembrane domain, helical stalk, and globular head (37). Three functions of the HN protein have been identified so far: 1) hemagglutinin activity for receptor binding, 2) neuraminidase activity for receptor cleavage by hydrolyzing sialic acid residues, and 3) fusion activation that interacts with the F protein (38). HN and F proteins work in tandem to infect the host cells. Binding of the HN protein to the sialic acid-containing receptors activates the F protein to fuse with the host cell membrane accompanied by structural rearrangement (39). Thus, communication between HN and F proteins is crucial for HPIV3 to mediate host cell infections (40).

Some mutations in the sialic acid receptor-binding sites of the HN protein have been reported to affect the physical strength of the HN-F interaction. The strains bearing the H552Q mutation in site II at the dimer interface are known to bind the receptor more avidly and activate the F protein more efficiently, with more extensive fusion *in vivo* (41, 42). In addition, the Q559R mutation has been shown to weaken the interaction between the HN dimers by inducing local conformational changes at the dimer interface (36). A single mutation that occurs at the dimer interface of the globular domain can affect the structure of the HN dimer and the features of the HN-F complex, thus altering the nature of HPIV3 infection in the host.

The Seoul HPIV3 strain showed some distinct amino acid substitutions, which were rarely observed in other reference strains. However, these sites did not correspond to the previously reported neutralization-related mouse monoclonal antibody-binding sites or active sites (37, 43). The G387S mutation was observed in all HPIV3 strains but was estimated to be one of the negative selection sites by FUBAR. This mutation was observed in one subcluster C3a strain reported in Argentina and in another study conducted on the long-term hospitalized patients (28, 44). The E514K mutation was observed in one reference strain isolated in Peru in 2007, which revealed neither positive nor negative selection. As both mutations have rarely been reported, the impact of these mutations on HPIV3 remains unclear.

The stalk region of the HN protein was previously noted at amino acid positions 54–136, and the globular head region at positions 137–572 (29). The F-specificity domain of the HN protein lies at amino acid positions 36–150 and 114–128 (45). The A22T and K31N mutations do not belong to the stalk, globular, or F-specific regions. Although we could not determine whether these mutations can significantly affect the infectivity of HPIV3 or alter the structure of proteins, further research is needed, as certain mutations have been reported to have a significant effect on viral activity (38).

Although HPIV3 infection has the second highest incidence after RSV infection, limited data on HPIV3 infections are available in Korea. Thus, in this study, a phylogenetic analysis was conducted following the sudden outbreak of HPIV3 infections, and new genetic lineages of the strain were identified. Continuous surveillance of HPIV3 is needed to detect novel mutations associated with antigen changes or the affinity characteristics of the virus.

In conclusion, the phylogenetic analysis using the *HN* gene showed that all Seoul HPIV3 strains were classified in subcluster C3; however, they formed a distinct group sharing the same rare mutations. This finding was in line with the genetic changes in other respiratory viruses after the huge aftermath of NPIs related to the COVID-19 pandemic. Thus, it is necessary to prepare in advance for unpredictable epidemics and magnitudes due to the rebound of respiratory pathogens. Although the impact of the emergence of this novel HPIV3 lineage remains unclear, this study has provided an opportunity to increase our understanding of the notable changes in HPIV3 during unseasonal epidemics.

## Acknowledgments

None.

## Data availability statement

The data that support the findings of this study are available from the corresponding author upon reasonable request.

---

**Figure S1.**
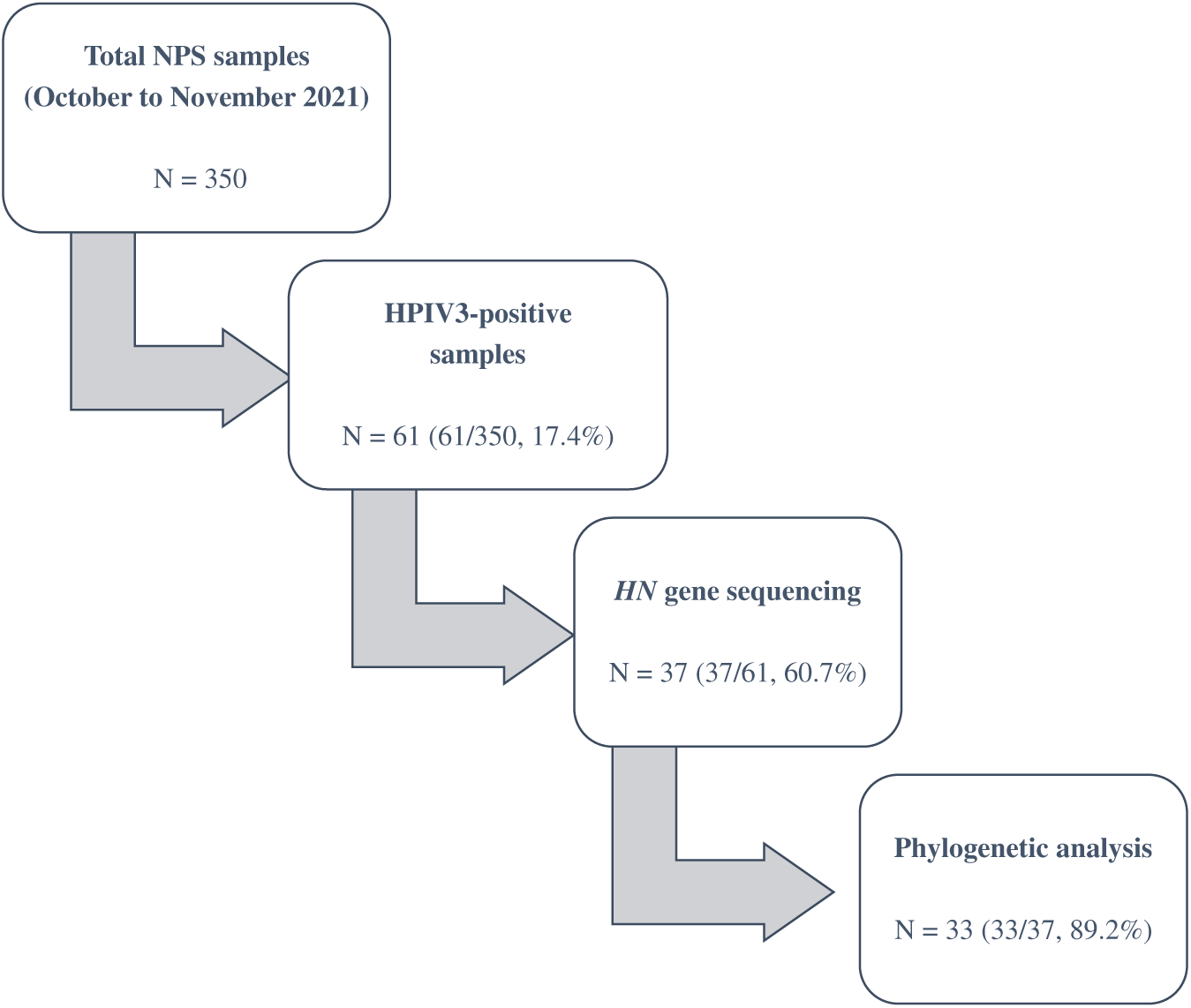
Schematic flow chart of sample number changes during the sample collection period (between October and November 2021).

